# Distributed cortical structural properties contribute to motor cortical excitability and inhibition

**DOI:** 10.1101/178301

**Authors:** Eran Dayan, Virginia López-Alonso, Sook-Lei Liew, Leonardo G. Cohen

## Abstract

The link between the local structure of the primary motor cortex and motor function has been well documented. However, motor function relies on a network of interconnected brain regions and the link between the structural properties characterizing these distributed brain networks and motor function remains poorly understood. Here, we examined whether distributed patterns of brain structure, extending beyond the primary motor cortex can help classify two forms of motor function: corticospinal excitability and intracortical inhibition. To this effect, we recorded high-resolution structural magnetic resonance imaging scans in 25 healthy volunteers. To measure corticospinal excitability and inhibition in the same volunteers we recorded motor evoked potentials (MEPs) elicited by single-pulse transcranial magnetic stimulation (TMS) and short-interval intracortical inhibition (SICI) in a separate session. Support vector machine (SVM) pattern classification was used to identify distributed multivoxel gray matter areas, which distinguished subjects who had lower and higher MEPs and SICIs. We found that MEP and SICI classification could be predicted based on a widely distributed, largely non-overlapping pattern of voxels in the frontal, parietal, temporal, occipital and cerebellar regions. Thus, structural properties distributed over the brain beyond the primary motor cortex relate to motor function.

## Introduction

Variation in local brain structure has been shown to be linked to performance in a range of motor functions (Kanai and Rees, 2011). Structure-function links of this sort were demonstrated at both the microstructural and macrostructural scales. For example, variation in microstructural white matter integrity in the body of the corpus callosum, as assessed with diffusion MRI is associated with variation in performance of a bimanual coordination task (Johansen-Berg et al., 2007). Similarly, individual differences in the macrostructural gray matter properties of the presupplementary motor area are linked to subjects’ ability to voluntarily select correct actions in the face of conflict (van Gaal et al., 2011). However, the link between the structural properties characterizing distributed brain networks and motor function remains incompletely understood.

In humans, transcranial magnetic stimulation (TMS) techniques have been vital in probing the physiological properties of the motor system (Dayan et al., 2013; Hallett, 2007; Rothwell, 1997). Two transcranial magnetic stimulation (TMS) protocols have been widely utilized as markers of motor corticospinal excitation and inhibition at rest. Motor evoked potentials (MEPs) elicited by single-pulse TMS over the primary motor cortex (M1) are a widely-used measure of instantaneous corticospinal excitability (Hallett, 2007; Rothwell, 1997; Rothwell et al., 1999). Similarly, short-interval intracortical inhibition (SICI), elicited by paired-pulse TMS over M1 is widely regarded as a measure of cortical inhibition (Kujirai et al., 1993; Rothwell et al., 2009). MEPs and SICIs relate with the structure of the primary motor cortex (Conde et al., 2012). However, since motor function does not rely solely on one cortical region (He et al., 2007; Matsumoto et al., 2006; Picard and Strick, 1996), we examined links between distributed structural properties of the cerebral cortex and MEPs and SICIs, an issue that has not been reported in the literature. We reasoned that since clear links between the structure or function of single brain regions and variability in subjects’ response to TMS were not identified to date, examining multivariate distributed substrates may provide an alternative approach. We thus evaluated whether differences in the magnitudes of MEPs and SICIs could be classified from subjects’ distributed whole-brain multi-voxel patterns of gray matter volume using whole brain machine learning pattern classification analysis.

## Methods

### Subjects

Data from 25 young, right-handed healthy volunteers (13/12 females/males; mean age 26.48± 5.15 STD) were used for analysis. Handedness was established using the Edinburgh Handedness Inventory (Oldfield, 1971). All subjects had unremarkable physical and neurological history, no MRI contradictions, and did not use any psychoactive medication. Written informed consent was obtained from all subjects prior to their participation in the study and all procedures were approved by the Combined Neuroscience Institutional Review Board, National Institutes of Health. All procedures were in accordance with approved guidelines.

### General procedure

All subjects underwent an imaging session and a stimulation session, which were administered separately (Fig 1A). The imaging session consisted of an anatomical scan (see details beneath). The stimulation sessions comprised single and paired-pulse TMS protocols, administered in an interleaved manner, where MEPs and SICIs were recorded respectively (Fig 1B). During the stimulation sessions, subjects were seated in a comfortable chair with their eyes open and were asked to stay relaxed and to not engage in conversation during the course of stimulation.

**Fig. 1.**
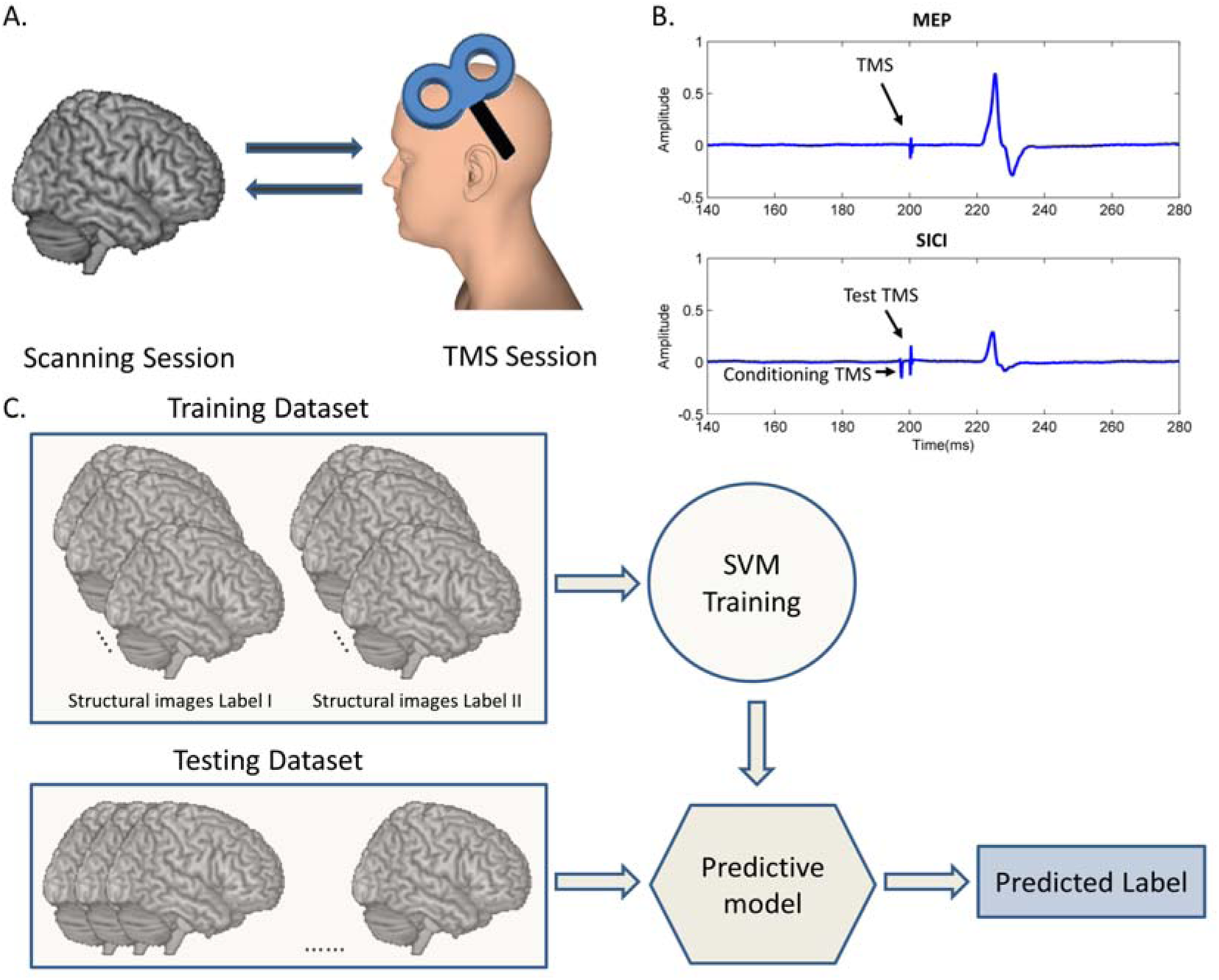
Design and analysis. (A) All subjects underwent a TMS session, where single and paired pulses were used to measures MEPs and SICIs and an imaging session where high resolution T1 weighted images were acquired, (B) MEPs were recorded from the FDI muscle following single TMS pulses delivered over primary motor cortex at 120% RMT. SICIs were recorded with a subthreshold conditioning stimulus at 80% RMT followed by a test stimulus at 120% RMT. (C) Pattern classification analysis was composed of two phases. In the training phase, a kernel SVM classifier was trained to distinguish between sets of anatomical images with known labels. A hyperplane that separates the training images according to their known labels was defined. Then, during the test phase, the performance of the classifier is tested on unlabeled images testing if the predictive model built during training can successfully classify the images. Fig 1A was drawn based on a human head model from: http://www.ir-ltd.net/. Used by Creative Commons license.

### EMG recordings

Electromyographic (EMG) traces were recorded via Ag/AgCl surface recording electrodes (7 mm x 4 mm recording area), placed over the right first dorsal interosseous (FDI) muscle. The active electrode was placed over the muscle belly and the reference electrode over the metacarpophalangeal joint of the index finger. Responses were acquired using a Neuropack MEB-2200 device (Nihon Kohden, Tokyo, Japan) through filters set at 10 Hz and 2 kHz with a sampling rate of 5 kHz, amplified (Micro-1401, Cambridge Electronic Devices, Cambridge, UK), and then recorded using the Signal software (Cambridge Electronic Devices, Cambridge, UK).

### TMS procedure

TMS was delivered through a figure-of-eight coil with an outer diameter of 70 mm (Magstim Co., Whitland, Dyfeld, UK) over the left motor cortex. The stimulators were triggered using the Signal software. The coil was held with the handle pointing backwards and laterally to evoke an anteriorly directed current in the brain (Sakai et al., 1997), and was optimally positioned to obtain MEPs in the FDI muscle. Using this configuration, single and paired pulses were delivered from a monophasic Magstim BiStim stimulator. We first localized the “motor hotspot” (defined as the point on the scalp at which single pulse TMS elicited MEPs of maximal amplitude from the right FDI). We then established each subject’s resting motor threshold (RMT), which corresponds to the minimum stimulation intensity over the motor hotspot, eliciting an MEP in the relaxed FDI of no less than 50 μV in 5 out of 10 trials.

Overall, the stimulation session included 40 stimulation trials, including 20 MEPs (at 120% RMT) and 20 SICI measures. MEPs and SICIs were administered in an interleaved and randomized manner, with an inter-trial interval of 5s, varying by up to 10%. SICIs were recorded as described previously (Kujirai et al., 1993), whereby a subthreshold conditioning stimulus (CS) at 80% of RMT preceded a test stimulus (120% RMT) by 3 ms.

### Imaging Setup

Imaging data was acquired with a 3.0-T GE Signa HDx scanner equipped with an 8-channel coil. High-resolution (1x1x1mm^3^) 3D magnetization prepared rapid gradient echo (MPRAGE) T1-weighted images were acquired (repetition time = 4.688 ms, echo time = 1.916 ms, slice thickness: 1.2 mm, slice spacing = 1.2 mm, acquisition matrix = 224x224 mm^2^, flip angle = 12°, 124 slices).

## Data Analysis

### TMS Data Analysis

Mean peak-to-peak amplitudes served as our primary outcome measure. Trial-to-trial variability in MEP amplitudes were additionally quantified using the coefficient of variation (CV), calculated across all MEP trials as follows:

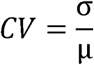

where σ denotes standard deviation and μ denotes the mean. For the SICI analysis, the mean peak-to-peak amplitude of the conditioned MEP was expressed as a percentage of the mean peak-to-peak amplitude of the unconditioned MEP. Trial-to-trial variation in SICI amplitudes was quantified using the CV metric, as described above. Parametric statistical tests were used to analyze MEP and SICI measurements after confirming the normality of the distributions using Kolmogorov-Smirnov tests. All tests were performed in SPSS 19 (Chicago, IL). Significance was set at alpha=0.05.

### Imaging data analysis

#### Image preprocessing

The VBM8 toolbox, part of Statistical Parametric Mapping 8, was used to preprocess subjects’ anatomical scans. The images were first normalized to the Montreal Neurological Institute (MNI) standard space and segmented for gray matter, white-matter and cerebro-spinal fluid (CSF) using the default segmentation routines of VBM8 (Gaussians per class 2,2,2,3,4,2; Bias regularization 0.0001; Bias FWHM 60mm cutoff; Affine regularization ICBM space template; Warping regularization 4; Sampling distance 3) and DARTEL normalization. The images were then subjected to pattern classification analysis, with the objective of finding distributed patterns of graymatter volume that could classify group differences in corticospinal excitability and inhibition at rest. This approach follows a recent body of research where multivariate morphometric parameters can be used to differentiate healthy controls from, for instance, patients with Alzheimer’s disease (Vemuri et al., 2008) or autism (Ecker et al., 2010b) based on machine learning pattern classification techniques (Bishop, 2006).

#### Pattern Classification

Pattern classification analysis was performed using the Pattern Recognition of Brain Image Data (PROBID) toolbox, on MATLAB 7. This analysis aimed to find patterns of gray matter volume that accurately classify group differences in MEPs and SICIs, treating subjects’ images as points in a high-dimensional space, corresponding to the number of voxels contained on each image (Dayan et al., 2014; Ecker et al., 2010b). Pattern classification analysis was used, rather than a mass-univariate approach, as it may potentially allow to detect more subtle multivariate structural substrates that contribute to variation in MEPs and SICIs.

The subjects were first split (using a median split) into groups, distinguishing between subjects for whom low and high mean MEP amplitudes and low and high SICIs were recorded (thus the 25 subjects were split into two groups of 12 subjects each, leaving out the median). The median split procedure enabled labeling of the data (into two groups in each classification), a prerequisite for supervised pattern classification. Modulated and normalized preprocessed gray matter images (see description of preprocessing steps, above) were then subjected to kernel support vector machine (SVM) classification (Boser et al., 1992). This procedure is composed of two phases. First, in the training phase, a kernel SVM classifier is trained to distinguish between modulated and normalized anatomical images, labeled according to the results of the median split analysis, described above. In this phase, a hyperplane that separates the images in the training dataset according to their known labels was defined. Then, during the test phase, the performance of the classifier was tested with a leave-two-out cross-validation procedure, whereby the test was administered n times (n = number of subjects), leaving a pair of subjects out on each iteration.

In this analytical framework *accuracy* also denotes the average between the classification’s *sensitivity* (proportion of subjects from class label I that were correctly classified as such) and *specificity* (proportion of subjects from class label II, who were correctly classified as such). As the input space used for classification was in voxel space, the weight vector normal to the hyperplane defined during training corresponds to the direction along which the images belonging to two groups differ the most. These inputs were then used to generate discrimination maps, which depicted a spatial map of the voxels that contributed the most to the discrimination among groups (Ecker et al., 2010b; Marquand et al., 2010; Mourão-Miranda et al., 2005). The maps depict voxels whose weights were at least 60% of the value of the voxel with the highest weight overall. This conservative threshold (Ecker et al., 2010a; Mourão-Miranda et al., 2005) allowed us to focus on the regions which most strongly discriminated among the groups. Discrimination maps were smoothed with a 3mm Gaussian kernel and cluster thresholded (10 voxels) for illustration purposes and the results were visualized using MRIcron (http://www.mccauslandcenter.sc.edu/mricro/mricron/).

Significance estimates for the accuracy of each classification were derived using a permutation test consisting of 5000 iterations. In this test, the classification procedure was repeated 5000 times, wherein labels were randomly assigned to subjects. In each permutation the cross-validation procedure was repeated and the number of times the accuracy levels exceeded those obtained with the original labeled data were counted, where p denotes the accumulated number of times divided by 5000.

## Results

Data from 25 young healthy volunteers were analyzed, testing the utility of using SVM classification of volumetric patterns of gray matter to predict group differences in corticospinal excitability and inhibition at rest, as measured with TMS-induced MEPs and SICs.

### Classification of MEP amplitudes

Subjects displayed substantial interindividual differences in mean peak-to-peak MEP amplitudes (Fig. 2A), which were normally distributed in this sample (Kolmogorov-Smirnov Z=0.162, p=0.091) Indeed, a median split of the data into two groups (n=12 each), of subjects who displayed low and high mean MEP amplitudes (henceforth, MEP_low_ and MEP_high_) resulted in significant differences between the groups (t_22_=5.234, p <0.0001), confirming the existence of sizable interindividual differences. These two groups did not differ in age (t_22_=0.922, p=0.366) or in their male/female distributions (Kolmogorov-Smirnov Z=1.021, P=0.249).

**Fig. 2.**
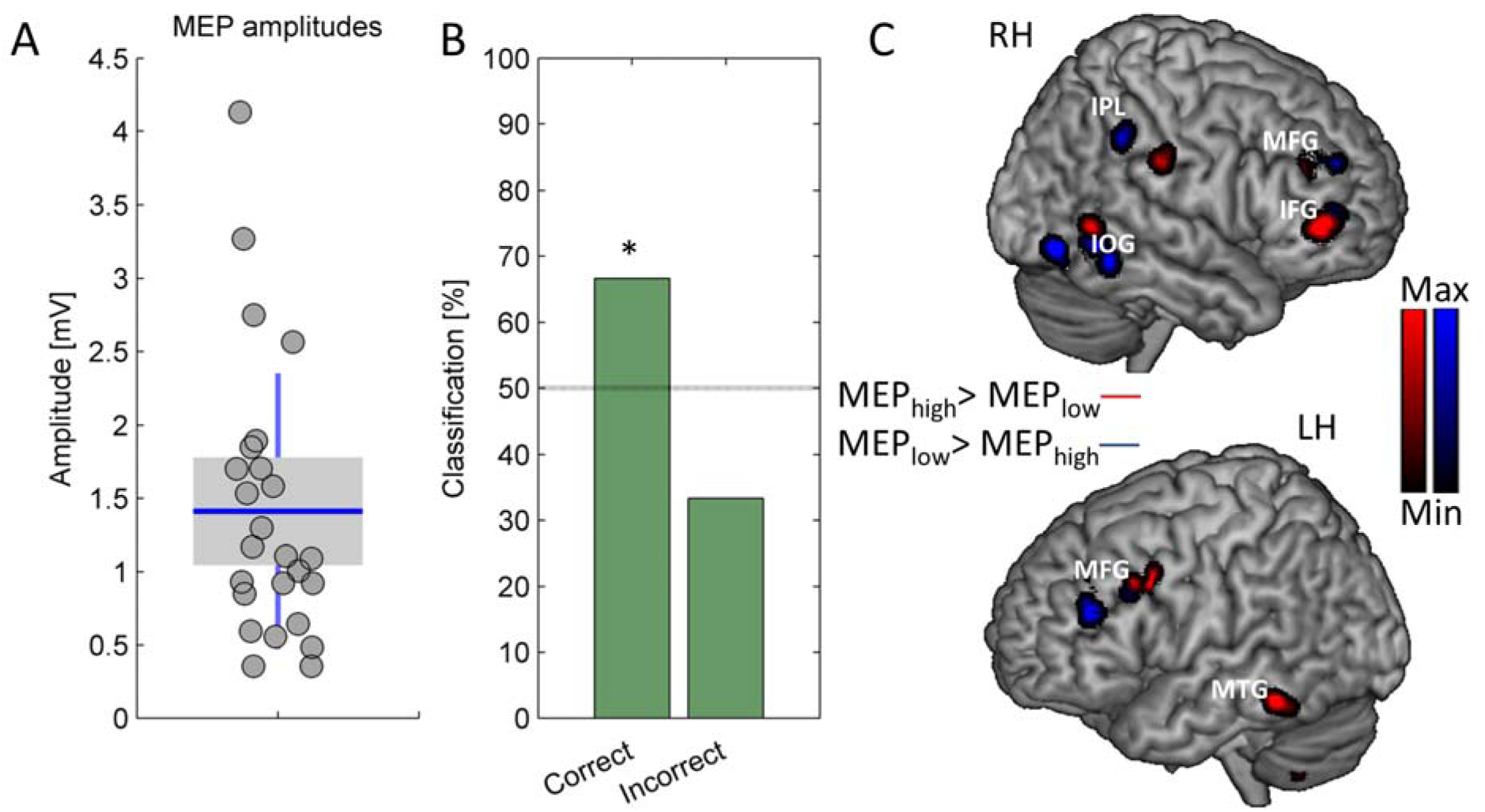
Pattern classification of MEP amplitudes. (A) Mean peak-to-peak MEPs differed by up to 168.4% among subjects. (B) Pattern classification of MEP amplitudes. Overall classification accuracy was significant at p <0.05, (random permutation test) (C) Discrimination maps depicting the weight of the voxels which contributed the most to the discrimination among the subjects who displayed low and high MEP amplitudes. Shown are regions where MEP_low_ > MEP_high_ (in blue) and where MEP_high_ > MEP_low_ (in red). LH, left hemisphere; RH, right hemisphere. IFG, inferior frontal gyrus; IOG, inferior occipital gyrus; IPL, inferior parietal lobule; MFG, middle frontal gyrus; MTG, middle temporal gyrus.

We first sought to test the degree to which group differences in MEP amplitudes could be predicted using whole-brain SVM classification. 75% of the subjects in the MEP_low_ group (the classification model’s ‘sensitivity’) and 58.33% of the subjects in the MEP_high_ group (the model’s ‘specificity’) were classified correctly, resulting in an overall accuracy of 66.67%, which was significantly better than chance (p <0.05, random permutation test) (Fig. 2B). Thus, these results reveal that patterns of gray matter allowed for a classification of group differences in MEP amplitudes. To more specifically identify the regions that contributed to this classification, discrimination maps were generated, depicting the weight of the voxels which contributed the most to the discrimination among the MEP_low_ and MEP_high_ groups (Fig. 2C, Table 1). The maps revealed that a widely distributed pattern of voxels composed of bilateral frontal and middle temporal, right inferior and anterior parietal and inferior occipital and left posterior cerebellar foci discriminated among the two groups.

**Table 1.** Brain regions that contributed the most to the discrimination among the subjects who displayed low and high MEP amplitudes. X,Y, Z coordinates (in MNI space) are displayed.

| Region | side | X | Y | Z |
| --- | --- | --- | --- | --- |
| ***MEP<sub>low</sub> > MEP<sub>high</sub>*** |  |  |  |  |
| Inferior Occipital Gyrus | R | 40.5 | -69 | -10.5 |
| Middle Frontal Gyrus | R | 22 | 37 | 29 |
| Middle Frontal Gyrus | L | -37.5 | 37.5 | 22.5 |
| Inferior Parietal Lobule | R | 52.5 | -39 | 40.5 |
| ***MEP<sub>high</sub> > MEP<sub>low</sub>*** |  |  |  |  |
| Posterior Cerebellum | L | -31.5 | -52.5 | -49.5 |
| Middle Temporal Gyrus | L | -55.5 | -40.5 | -15 |
| Middle Temporal Gyrus | R | 56 | -52.5 | 3 |
| Inferior Frontal Gyrus | R | 37.5 | 45 | 3 |
| Middle Frontal Gyrus | R | 27 | 37 | 25 |
| Middle Frontal Gyrus | L | -37.5 | 10.5 | 36 |
| Postcentral Gyri | R | 58.5 | -22.5 | 33 |

We next quantified the trial-to-trial variability in amplitudes with the CV statistic (Fig. 3A). This allowed us to assess the contribution of more transient and unspecific factors to the classification of MEPs (for instance, movement of the TMS coil along the stimulation site, slight changes in the orientation of stimulation, etc). MEP CVs differed substantially among subjects and were insignificantly correlated with mean MEP amplitudes (r=-0.268, p=0.195), suggesting that these two measures were largely independent of one another in the current sample of subjects. A median split of the data into two groups (n=12 each) of subjects who displayed high and low CVs (henceforth, MEPCV_low_ and MEPCV_high_) resulted in significant difference between the two groups (t_22_=5.53, p <0.0001). These two groups did not differ in age (t_22_=0.269, p=0.791) or in their male/female distribution, which were identical.

**Fig. 3.**
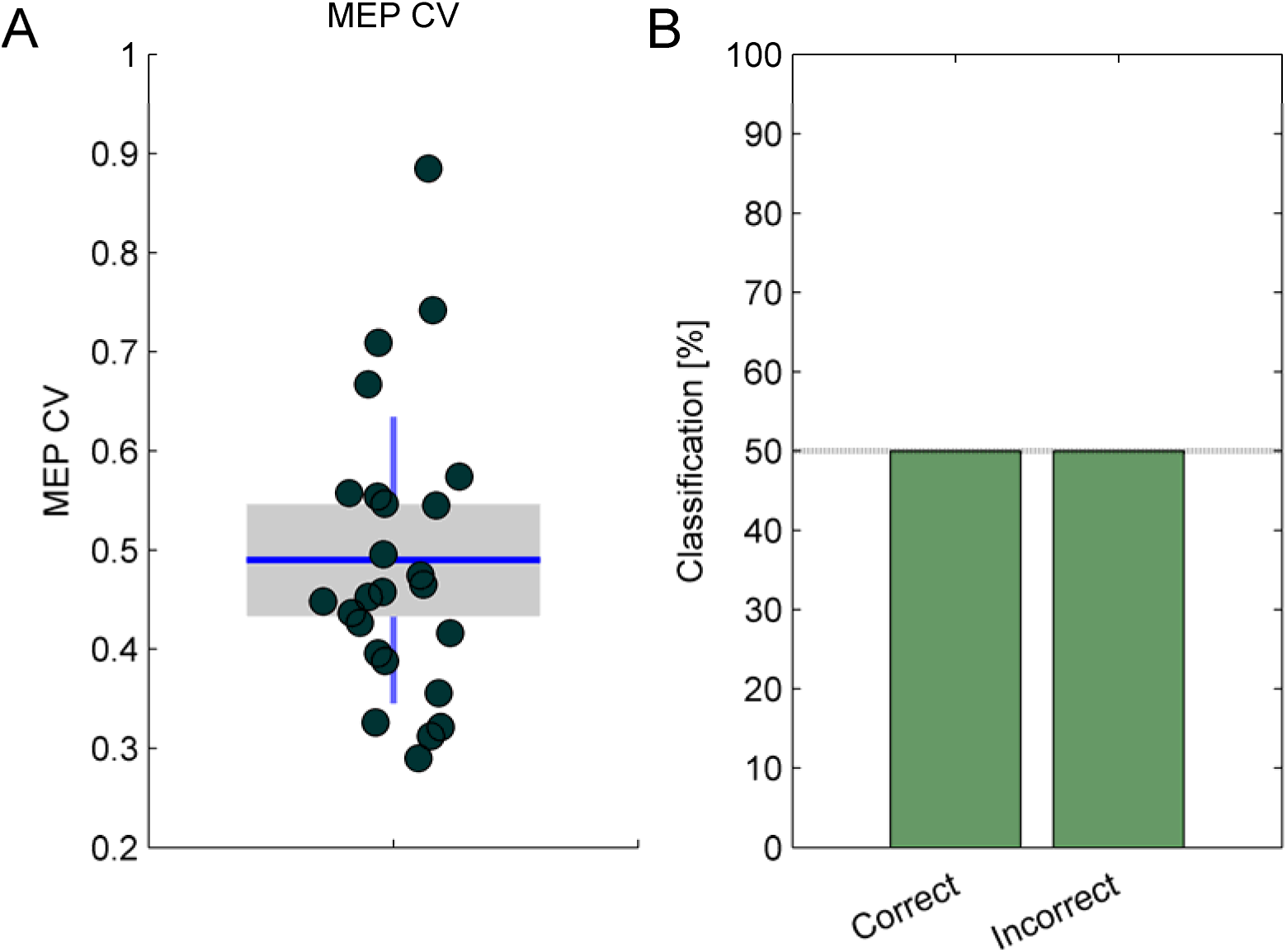
Pattern classification of MEP trial-to-trial variation. (A) Trial-to-trial variability in MEP amplitudes, quantified with CV statistic, differed among subjects by up to 101.3% (B) Overall classification accuracy was at chance levels (50%)

The trial-to-trial variation in MEP amplitudes could not be accurately classified in relation to gray matter volumetric patterns. Only 41.67% of the subjects in the MEPCV_low_ (sensitivity) and 58.33% of the subjects in the MEPCV_high_ (specificity) groups were classified correctly, resulting in an overall accuracy of 50% (Fig. 3B). The accuracy of this classification, which did not differ from chance levels, was not statistically significant (p= 0.58)

### Classification of SICI amplitudes

SICI amplitudes were also variable among subjects (Fig. 4A) and the data was also normally distributed in this sample (Kolmogorov-Smirnov Z=0.126, p=0.2). Indeed, a median split of the data into two groups (n=12 each) of subjects who displayed low and high mean SICIs (henceforth, SICI_low_ and SICI_high_) resulted in significant differences between the groups (t_22_=6.48, p <0.0001). These two groups did not differ in age (t_22_=0.154, p=0.879) or in their male/female distributions (Kolmogorov-Smirnov Z=0.408, P=0.996).

**Fig. 4.**
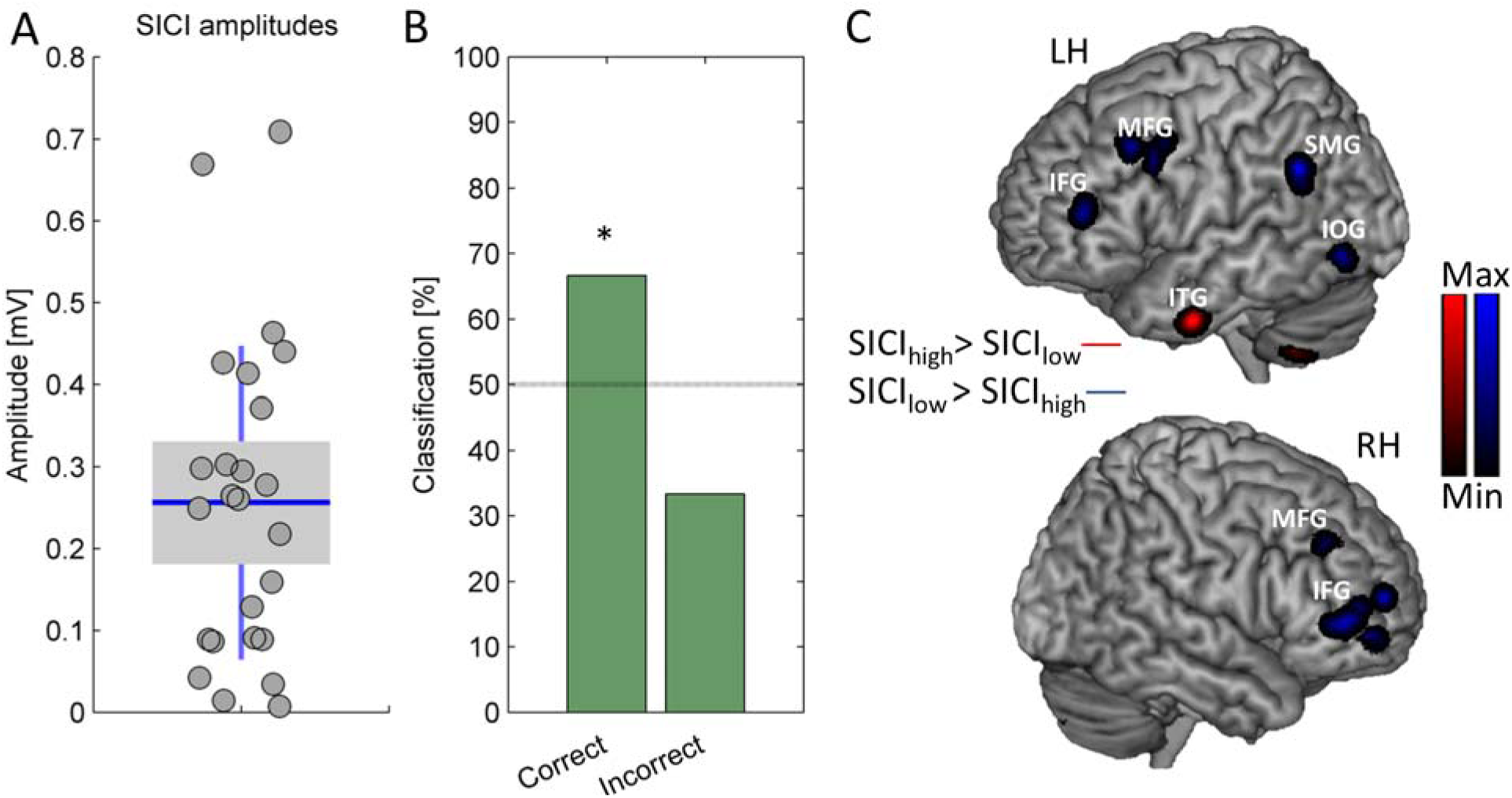
Pattern classification of SICI amplitudes. (A) Mean peak-to-peak SICIs differed by up to 195.83% among subjects. (B) Overall classification accuracy was significant at p <0.05, (random permutation test) C. Discrimination maps depicting the weight of the voxels which contributed the most to the discrimination among the subjects who displayed low and high MEP amplitudes. Shown are regions where SICI_ow_ > SICI_high_ (in blue) and where SICI_high_ > SICI_low_ (in red). LH, left hemisphere; RH, right hemisphere. IFG, inferior frontal gyrus; IOG, inferior occipital gyrus; ITG, inferior temporal; gyrus MFG, middle frontal gyrus; SMP, supramaginal gyrus.

We next tested whether SICI mean amplitudes could be predicted using whole brain SVM classification. 58.33% of the subjects in the SICI_low_ group (the model’s sensitivity) and 75% of the subjects in the SICI_high_ group (the model’s specificity) could be classified correctly (Fig 6A), which together summed up to 66.67% accuracy, significantly better than chance levels (p <0.05, random permutation test; Fig 4B). Discrimination maps were then generated in order to identify the regions which contributed to this classification. A distributed pattern of voxels in bilateral frontal and posterior cerebellar and left inferior occipital and inferior parietal foci contributed mostly to the classification of group differences in SICI amplitudes (Fig 4C, Table 2).

**Table 2.** Brain regions that contributed the most to the discrimination among the subjects who displayed low and high SICI amplitudes. X,Y, Z coordinates (in MNI space) are displayed.

| **Region** | **side** | **X** | **Y** | **Z** |
| --- | --- | --- | --- | --- |
| ***SICl<sub>low</sub> > SICl<sub>high</sub>*** |  |  |  |  |
| Inferior Occipital Gyrus | L | -39 | -70.5 | -7.5 |
| Inferior Frontal Gyrus | R | 39 | 45 | 3 |
| Inferior Frontal Gyrus | L | -37 | 42 | 13 |
| Supramarginal Gyrus | L | -51 | -51 | 31.5 |
| Middle Frontal Gyrus | R | 25 | 34 | 35 |
| Middle Frontal Gyrus | L | -34 | 19.5 | 40.5 |
| ***SICl<sub>high</sub> > SICl<sub>low</sub>*** |  |  |  |  |
| Posterior Cerebellum | L | -31.5 | -51 | -49.5 |
|  | R | 12 | -84 | -37.5 |
| Inferior Temporal Gyrus | L | -46.5 | -4.5 | -33 |

The trial-to-trial variability in SICIs, quantified with the CV statistic (Fig. 5A), differed substantially among subjects and was significantly inversely correlated with mean SICIs (r=-0.498, p<0.02). A median split of this data into two groups (n=12 each) of subjects who displayed high and low SICI CVs (henceforth, SICICV_low_ and SICICV_high_) resulted in significant difference between the two groups (t_22_=5.818, p <0.0001). These two groups did not differ in age (t_22_=0.4, p=0.969) or in their male/female distributions (Kolmogorov-Smirnov Z=0.408, P=0.996).

**Fig. 5.**
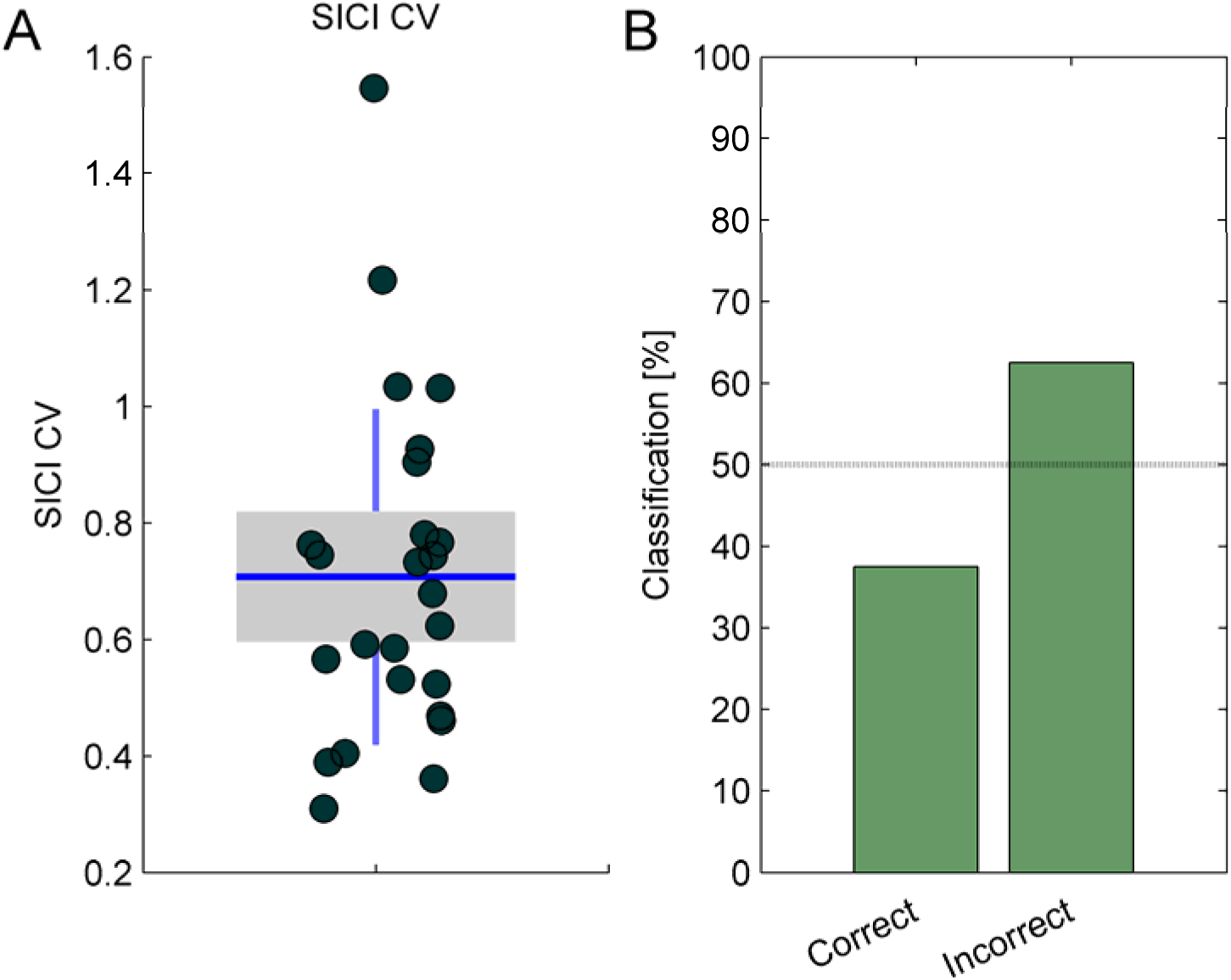
Pattern classification of SICI trial-to-trial variation. (A) Trial-to-trial variability in SICI amplitudes, quantified with CV statistic, differed among subjects by up to 133.3% (B) Overall classification accuracy was worse than chance levels.

**Fig. 6.**
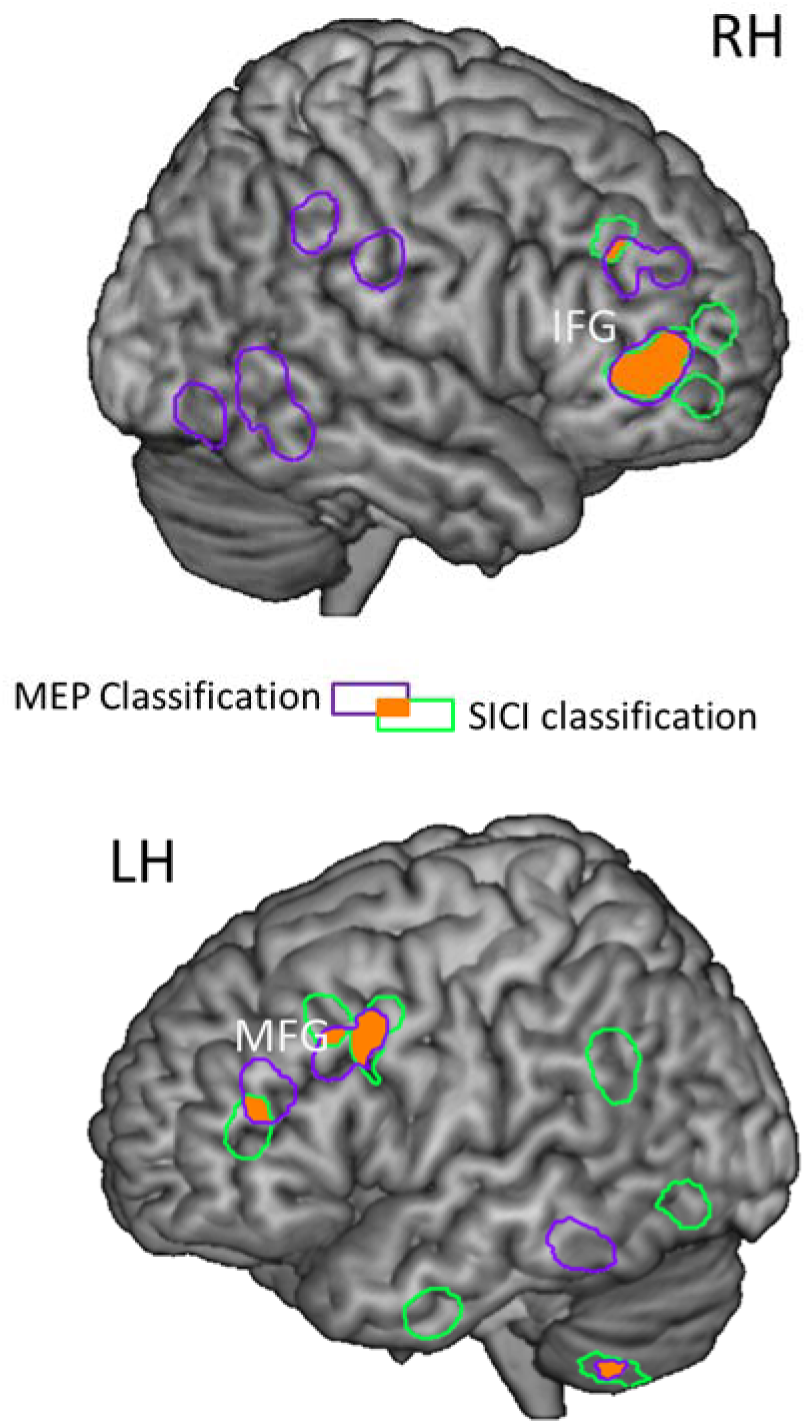
Similarities in the classification of MEP and SICI amplitudes. Overlaying the discrimination maps identified in the classification of each of these measures (depicted in purple and green contours) revealed that apart from a partial overlap in bilateral frontal cortex and right cerebellum (overlap in orange) the maps were mostly non-overlapping. IFG, inferior frontal gyrus; MFG, middle frontal gyrus.

We next assessed the degree to which group differences in the trial-to-trial variability of SICIs could be predicted using whole brain SVM classification. 41.67% of the subjects in the SICICV_low_ group (sensitivity) and 33.33% of the subjects in the SICICV_high_ group (specificity) were classified correctly. Combined, whole brain SVM of SICI CVs thus resulted in 37.5% accurate classification, a proportion which was not statistically significant (p=0.925; Fig. 5B).

### Similarities between MEP and SICI classification

Finally, we assessed similarities in the classification of MEPs and SICIs, as our results indicate that both were statically significant. In the current dataset MEPs and SICIs were insignificantly correlated within subjects (r=-0.169. p= 0.419). Overlaying the discrimination maps identified in the classification of each of these measures (Fig. 2C and Fig. 4C) revealed that the maps were largely non-overlapping (Fig. 6).

## Discussion

We tested the feasibility of classifying group differences in mean MEP and SICI, elicited by single and paired-pulse stimulation of M1, from multi-voxel patterns of gray matter volume, aiming to find links between distributed structural properties and motor function. Mean MEP classification could be achieved with significant accuracy levels from a widely distributed pattern of voxels composed of regions in frontal, parietal, temporal occipital and cerebellar regions. Similarly, mean SICI was predicted, with similar accuracy levels, from a widely distributed pattern of voxels which included foci in frontal, parietal, temporal and occipital cortices, as well as the cerebellum. The prediction was specific to mean MEP peak-to-peak amplitudes and SICIs. Group differences in the trial-to-trial variation in neither MEPs nor SICIs could be classified from patterns of gray matter volume.

Our results reveal that distributed patterns of gray matter volume, extending well beyond M1 allowed for an accurate group classification of MEP and SICI amplitudes. While MEPs are widely used for probing the physiology of M1, multiple lines of evidence suggest that they do not reflect a simple read-out of neuronal processes occurring within M1, but rather also tap into physiological processes that occur outside of this structure (Bestmann and Krakauer, 2015). This is strongly demonstrated by dual-site stimulation paradigms where a conditioning TMS pulse is delivered to various cortical regions prior to a test stimulus in M1 (Dayan et al., 2013), establishing the functional connectivity of these regions with M1, while revealing the influence these regions may exert over MEPs (Bestmann and Krakauer, 2015; Liew et al., 2014). Along these lines, several of the regions which were found here as implicated in the classification of MEP and SICI have been shown to interact with M1 based on dual-site TMS stimulation. For example, connectivity between M1 and ventral premotor (Buch et al., 2010) and dorsal premotor (O’Shea et al., 2007) cortices has been demonstrated, which fits well with the dense inputs between these regions and M1 in the monkey (Dum and Strick, 2005; Hoshi and Tanji, 2007). Similarly, connectivity between the cerebellum and M1 has been demonstrated with dual-site TMS (Daskalakis et al., 2004), consistent with cerebellar –motor cortex loops found in the monkey (Kelly and Strick, 2003). Altogether, using a whole brain analysis and a nonrestricted and unbiased pattern classification method we have not found multivoxel patterns in M1. In summary, our results identify a relationship between distributed multi-voxel patterns of brain volume in extra-motor cortical regions and variability in MEP and SICI.

In addition to our focus on group classification of differences in corticospinal excitability and inhibition, which we studied by quantifying mean MEPs amplitudes and SICIs, we also tested whether there were differences in the trial-to-trial variability subjects display, quantified with the CV statistic. This allowed us to assess the contribution of more transient and unspecific factors to the classification of MEPs and SICIs. Contrary to mean MEP and SICI amplitudes, intra-subject variation could not be classified from patterns of gray matter volume. It may thus be that intra-subject variation is indeed induced by more transient and state-dependent factors, which were not controlled for in this study, whereas group differences in mean MEP and SICI amplitudes were driven by more stable, possibly pre-existing state-independent factors.

Inter- and intraindividual differences in MEP and SICI amplitudes have been widely reported before, and were ascribed largely to transient, spontaneous and state-dependent factors. Variation in MEP measurement, for instance, has been attributed to spontaneous fluctuations in corticospinal and segmental motoneuron excitability (Kiers et al., 1993). Likewise, various transient physiological states such as pre-stimulation muscle activation (Darling et al., 2006), central (Temesi et al., 2014) and more localized (muscle-specific) fatigue (Taylor and Gandevia, 2001), response preparation (Mars et al., 2007) and attention (Rosenkranz and Rothwell, 2004; Thomson et al., 2008) modulate MEPs and SICIs. Our results cannot exclude the contribution of transient or spontaneous factors to variability in MEPs and SICIs. However, our data suggests that stable, possibly pre-existing non state-dependent neuroanatomical substrates may allow for the classification of group differences in corticospinal excitability and inhibition. Thus, intrinsic (Goetz et al., 2014) and stable properties like brain structure could contribute to the variability in corticospinal excitability and inhibition.

Non-transient factors may also contribute to the variability in MEPs or SICIs. For instance, age and sex interact with the trial-to-trial variability of MEP amplitude (Pitcher et al., 2003), but in our current results, none of the classified groups differed in age or in male/female ratios, so the contribution of these factors to the results reported here is unlikely. Similarly, the physical parameters of stimulation may also contribute to the variability in MEPs or SICIs, but their contribution to their variability is not trivial. For instance, the relationship between mean MEP amplitude and stimulation intensity is not necessarily linear (Darling et al., 2006). Likewise, variability in MEPs does not decrease when more elaborate methods are used to localize M1, such as stereotactic neuronavigation (Jung et al., 2010). The contribution of factors such as stimulation intensity and fluctuations in coil positioning to the degree to which variation in MEPs and SICIs relates to brain structure remains to be tested in future research.

As subjects’ attention has been found to influence MEPs (Mars et al., 2007) and SICIs (Rosenkranz and Rothwell, 2004; Thomson et al., 2008), one possibility that warrants consideration is that the structural differences between subjects who showed higher and lower MEPs and SICIs may relate to interindividual differences in attention, which in turn influenced MEP amplitudes and SICIs. Regions such as the dorsolateral prefrontal cortex (dlPFC), inferior frontal gyrus, supramarginal gyrus and middle temporal gyrus, found here as implicated in the classification of MEP amplitudes and SICIs are considered to be a part of the dorsal and ventral attention networks (Fox et al., 2006; Vossel et al., 2014). Moreover, correlations between regional structural properties of several of the brain regions found here as implicated in the classification of MEP and SICI with attention or attention-related functions have been reported before (Smolker et al., 2015; Westlye et al., 2010). For instance, significant associations were found between the executive component of attention and cortical thickness in the middle and superior temporal gyri, inferior frontal gyrus and dlPFC, and reduced gray matter volume in dlPFC is associated with better performance in the monitoring and updating of working memory (Smolker et al., 2015), functions which require attentional control (Fougnie, 2008). Future research may explore the relationship between attention-related regions and corticospinal excitability.

Although several previous reports found a relationship between SICI and MEP estimates (Roshan et al., 2003; Sanger et al., 2001), a systematic examination revealed that SICIs is dependent on the intensity of the test TMS pulse, rather than the size of the test MEP per se (Garry and Thomson, 2009). These results are consistent with our findings that MEP and SICI amplitudes did not covary among subjects and the regions which contributed to classification of differences along these measures were largely independent. Still, because of the lack of covariation in MEPs and SICIs among our sample of subjects, the classification analysis for these two measures was based on different subject-groupings. Thus, although our results suggest that differences in MEPs and SICI amplitudes may possibly originate from variation in largely non-overlapping structural substrates additional data is needed in order to confirm this suggestion.

Our goal here was to establish the feasibility of classifying group differences in MEP and SICI amplitudes based on multi-voxel patterns of brain volume, aiming to find links between distributed structural properties and motor function. An advantage afforded by this approach is that it enables us to detect subtle and distributed morphological differences between subjects possibly masked by a mass-univariate approach such as voxel-based morphometry (Dayan et al., 2014; Ecker et al., 2010b). However, the supervised learning approach we used here required splitting of the dataset into groups which thus reduced the true variation of the dataset. Links between multivariate representations of brain structure and a more continuous variation in MEP and SICIs should be established in future studies.

## Acknowledgments

This work was supported by the Intramural Research Program of the National Institute of Neurological Disorders and Stroke, National Institutes of Health. Virginia López-Alonso was supported by an FPU fellowship from Ministerio de Educación, Cultura y Deporte, Spain. The study utilized the high-performance computational capabilities of the Biowulf Linux cluster at the National Institutes of Health, Bethesda, Md. (http://biowulf.nih.gov). We thank Ryan Thompson for assistance in the preparation of this manuscript.

## Author Contributions

All authors designed the study; V.L.A and S.L.L performed the experiments. E.D and V.L.A analyzed the data. All authors wrote and reviewed the manuscript.

